# Engineering the *C. elegans* genome with a nested, self-excising selection cassette

**DOI:** 10.1101/2025.05.01.651742

**Authors:** Theresa V Gibney, Ariel M Pani

## Abstract

*C. elegans* is a powerful model for dissecting biological processes *in vivo*. In particular, the ease of generating targeted knock-in alleles makes it possible to visualize and functionally modify endogenous proteins to gain fundamental insights into biological mechanisms. Methods for *C. elegans* genome engineering typically utilize selectable markers, visual screening for fluorescence, or PCR genotyping to identify successfully edited animals. A common genetic tool known as the Self-Excising Cassette (SEC) combines drug and phenotypic selection, which makes it possible to screen large numbers of progeny rapidly and with minimal hands-on effort. However, N-terminal and internal knock-ins using the SEC cause loss of function until the selectable marker cassette is excised, which makes it impossible to isolate homozygous lines for essential genes prior to SEC excision. To simplify generating knock-ins for essential genes, we developed a Nested, Self-Excising selection Cassette (NSEC) that is located entirely within a synthetic intron and does not interfere with the expression of endogenous, N-terminally-tagged NSEC-fusion proteins. This innovation makes it possible to isolate homozygous lines for N-terminally tagged genes prior to selectable marker excision and allows for a standardized workflow to generate N-terminal and internal tags in any background and without the need for genetic balancers. We designed versions of NSEC that include an optional auxin-inducible degron tag and mTurquoise2, GFP, mStayGold, mNeonGreen, or mScarlet-I fluorescent proteins for experimental flexibility. The NSEC expands our molecular toolbox and enhances the scalability, efficiency, and versatility of *C. elegans* genome engineering.

## Introduction

*C. elegans* is a genetically tractable model organism used to investigate a broad range of fundamental biological processes. Its small size, optical transparency, and relative ease of genome engineering compared to other multicellular organisms make *C. elegans* a particularly powerful system for studies that use *in vivo* imaging. Efficient genome engineering approaches based on CRISPR/Cas9 have revolutionized genetic manipulations in *C. elegans*, providing researchers with unprecedented abilities to generate precise fluorescent protein knock-ins and other modifications at endogenous loci (reviewed by DICKINSON AND GOLDSTEIN 2016). When tagging endogenous genes, it is critical to choose tag locations that do not affect the function of the resulting fusion protein. Many proteins do not tolerate C-terminal tags because specific C-terminal sequences are often required for proper localization, function, or post-translational modification (CLARKE 1992; CHOY *et al*. 1999; SNAPP 2005; ROBERTS *et al*. 2008). Both the N- and C-termini are required for native properties of some proteins, and internal knock-ins may be needed to preserve their functions (WALL *et al*. 1995; ADJOBO-HERMANS *et al*. 2011; ARMENTI *et al*. 2014; BENDEZU *et al*. 2015). For some genes, the presence of alternative splice forms may also necessitate using an N-terminal or internal tag (KEELEY *et al*. 2020). In other cases, specific tag locations may be needed due to native protein cleavage, and/or there may be experimental requirements to tag the protein in multiple locations (SOHR *et al*. 2019). Accordingly, there is a need for rapid, flexible, and efficient methods that can be used to tag proteins in any location.

Tagging endogenous genes in *C. elegans* typically relies on homology-directed repair (HDR) to insert a repair template with site-specific homology arms at the site of a Cas9-induced double-stranded DNA break. Multiple techniques leverage Cas9-triggered HDR to tag endogenous genes in *C. elegans* (reviewed by DICKINSON AND GOLDSTEIN 2016). Relatively small sequences (<140bp) can be inserted using a synthetic, single-stranded DNA oligonucleotide repair template with short homology arms (PAIX *et al*. 2014; ZHAO *et al*. 2014; WARD 2015). Larger sequences, up to the size of a fluorescent protein, can also be inserted using PCR-generated double-stranded DNA repair templates with short homology arms (PAIX *et al*. 2015; GHANTA AND MELLO 2020), but with lower efficiency than smaller inserts. While these approaches are effective for some sites, success rates are variable, and post-injection screening can be highly labor-intensive. Screening is particularly time-consuming for loci with low editing efficiency and/or knock-ins targeting genes that are expressed at levels too low to visualize on a fluorescence stereomicroscope. To simplify the process of identifying rare knock-ins, several strategies incorporate selectable markers encoded in the homologous repair template (DICKINSON *et al*. 2013; ARMENTI *et al*. 2014; DICKINSON *et al*. 2015) that make it possible to screen for successful knock-ins with minimal hands-on effort and a flexible timeline. A common method uses a Self-Excising selection Cassette (SEC) that includes a hygromycin resistance gene, a dominant *sqt-1(d)* marker that confers a visible rolling phenotype, and heat-shock-driven *cre* recombinase (DICKINSON *et al*. 2015). Unlike selection methods that rely on rescuing a mutant, the SEC has the key advantage that it can be used in any genetic background. In addition to streamlining the process of identifying knock-in animals, the roller phenotype simplifies subsequent strain crossing by serving as a visible proxy for the knock-in genotype. The SEC is flanked by loxP sites and can be excised from the genome by Cre-lox recombination.

Despite the ease of screening for knock-ins, the original SEC has design features that complicate tagging many genes of interest. Knocking the SEC into an endogenous gene interferes with the expression of downstream coding and/or untranslated regions (UTRs) prior to SEC excision (DICKINSON *et al*. 2015). Until the SEC is excised, N-terminal SEC insertions typically act as strong loss-of-function alleles while internal knock-ins truncate the endogenous protein. Therefore, N-terminal and internal SEC knock-in alleles for essential genes are typically not viable as homozygotes, and lines must be maintained prior to SEC excision by picking rolling worms in each generation or by crossing in a genetic balancer. C-terminal SEC knock-ins do not disrupt the coding sequence but result in a transcript where the endogenous 3’ UTR is replaced with a *let-858* 3’ UTR located in the SEC prior to excision. While many genes tolerate this 3’ UTR replacement, essential genes with critical regulatory elements in the 3’UTR (THOMPSON *et al*. 2006; MERRITT *et al*. 2008; OLDENBROEK *et al*. 2013) may not. The ease of balancing SEC alleles depends on the availability of balancer strains for the targeted region and their genetic and phenotypic compatibility with the knock-in’s parental strain, which may include mutations or insertions at multiple loci. Although it is possible to excise the SEC from unbalanced heterozygous lines, identifying excised progeny can be very challenging because their movement phenotype is indistinguishable from the 25% of progeny that do not carry a knock-in allele. As a result, it is not possible to use a standardized workflow for making N-terminal and internal tags on essential genes using the SEC, and each knock-in requires individualized efforts that limit throughput and can become an impediment to progress.

We sought to reengineer the SEC to allow for functional N-terminal and internal knock-ins prior to cassette excision while maintaining the core advantages of the original design. To do so, we designed a Nested, Self-Excising selection Cassette (NSEC) that is entirely embedded within a synthetic intron and does not interfere with expression or function of N-terminal and internal NSEC knock-in alleles. To validate this approach, we endogenously tagged the essential *ERK1/2* homolog *mpk-1* at its N-terminus with mNeonGreen(mNG) and an auxin-inducible degron (AID) using the SEC or NSEC. As predicted, the *mNG^SEC^3xFlag::AID::mpk-1* knock-in (^ denotes an artificial intron prior to SEC/NSEC excision) caused loss of MPK-1 function, and knock-in lines could only be maintained as heterozygotes. In contrast, an otherwise identical NSEC-based knock-in preserved endogenous MPK-1 function. *mNG::AID^NSEC^mpk-1* animals were viable as homozygotes, did not display loss-of-function phenotypes, and mNG::AID^NSEC^MPK-1 protein was localized correctly. To facilitate wider use in the *C. elegans* community, we also generated ten NSEC plasmid backbones that can be used to clone homologous repair templates with the fluorescent proteins mTurquoise2, GFP, mStayGold, mNG, and mScarlet-I, with or without an AID tag. NSEC provides a scalable and user-friendly approach for tagging endogenous genes in any location without initially disrupting their function, which simplifies the workflow to generate knock-ins and has potential to facilitate large-scale endogenous tagging efforts.

## Results/Discussion

To prevent N-terminal and internal SEC knock-ins from disrupting gene function, we sought to redesign the SEC so that it could be embedded entirely within an artificial intron. We reasoned that it should be possible to hide the SEC within an intron by flanking the selectable marker cassette with splice donor and acceptor sites and eliminating transcriptional terminators and other splice acceptors in the same orientation as the gene of interest. We first removed a splice acceptor sequence, HA tag, and *let-858* 3’UTR that are present at the 5’ end of the original SEC. We then rearranged the *HygR, sqt-1(d)*, and *hs>cre* genes that make up the SEC so that all three are transcribed in the opposite orientation as the tagged gene of interest (Figure 1). For an intronic SEC to be functionally “hidden” from an endogenously tagged gene, it should also be essential to remove all splice acceptors within the selectable marker genes to ensure that the entire cassette is spliced out of the target gene’s mRNA. To identify splice acceptors within the rearranged SEC, we used NetGene 2 - 2.42 (BRUNAK *et al*. 1991; HEBSGAARD *et al*. 1996) to produce neural network predictions of *C. elegans* splice sites in our modified sequence. For predicted splice acceptors in coding sequences, we made synonymous substitutions to disrupt the splice acceptor function without altering the encoded protein. For predicted splice acceptors in non-coding sequences, we made semi-random single nucleotide substitutions or added a single nucleotide to disrupt the splice acceptor function (Supplemental Note 1A). To reduce the overall size of the selectable marker cassette, we removed extraneous sequence between the *rps-0* promoter and *hygromycin phosphotransferase* and replaced the *unc-54* 3’UTR in the *HygR* gene with a 107 bp minimal *let-858* 3’UTR. We also shortened the *tbb-2* 3’ UTR in the *hs>cre* gene to 40 bp. We flanked this redesigned SEC with loxP sites and nested it within a synthetic intron that included unique intronic sequences to facilitate subsequent cloning by Gibson assembly (Figure 1; Supplemental Note 1). We named this modified selectable marker cassette the Nested Self-Excising selection Cassette (NSEC). To generate a user-friendly backbone plasmid for cloning homologous repair templates, we added mNG::AID along with extended flexible linker sequences and restriction sites located between functional sequence features (Figure 1; Supplemental Note 1).

**Figure 1.**
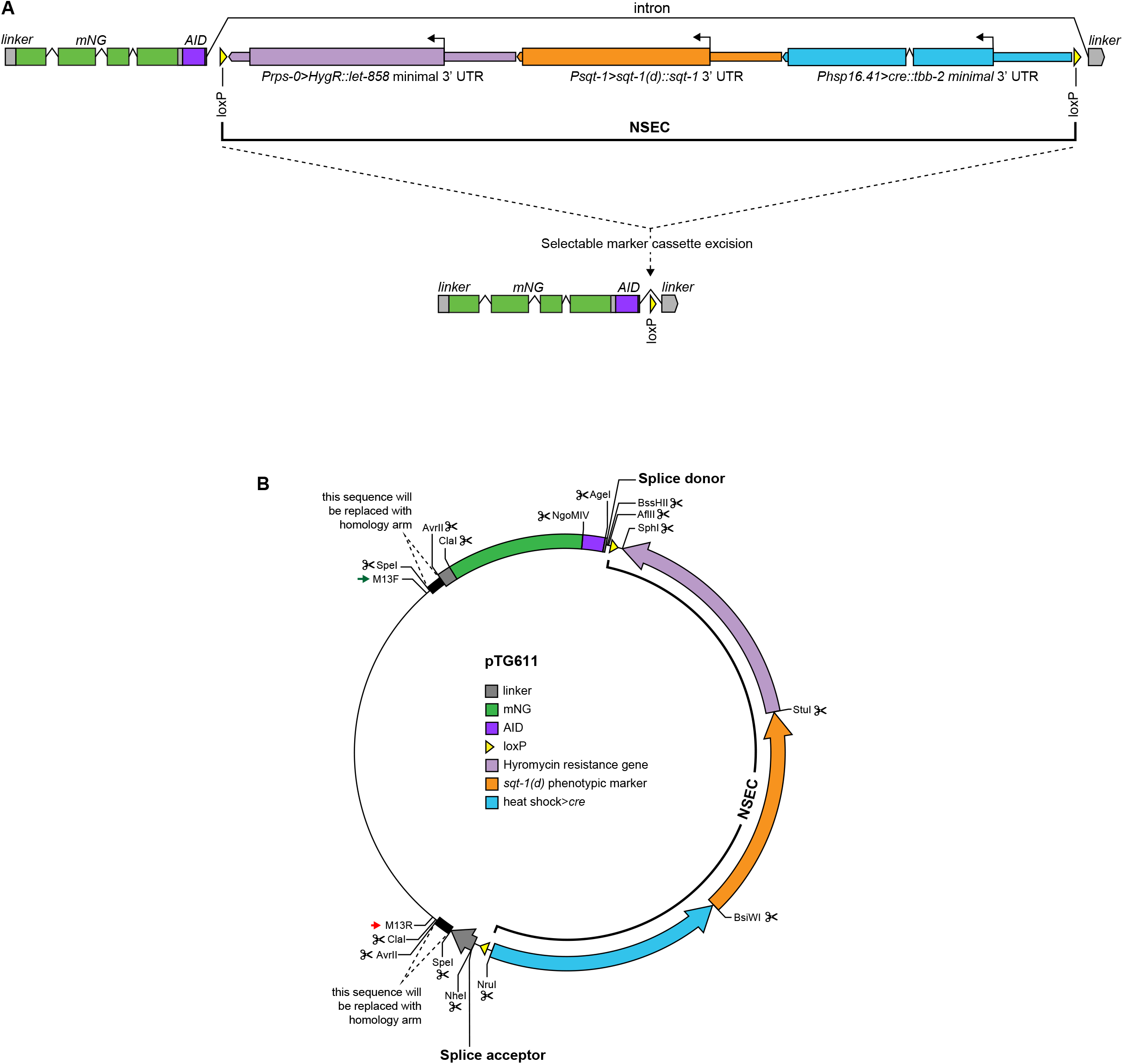
Design of the Nested, Self-Excising selection Cassette.

To assess performance of the NSEC for N-terminal knock-ins, we decided to endogenously tag an essential gene with a known localization pattern. As a test case, we chose the *ERK1/2* homolog *mpk-1*, which is essential for larval development and adult fertility (reviewed by SUNDARAM 2013). Endogenously tagged MPK-1 localizes to the nucleus in a subset of cells where MAPK/ERK is active (RASMUSSEN AND REINER 2021), and its subcellular localization is clearly distinguishable from the uniform fluorescence observed in N-terminal knock-ins with the original SEC prior to excision (DICKINSON *et al*. 2015). To assess NSEC functionality, we generated knock-ins tagging *mpk-1* at its N-terminus using either the original SEC or NSEC for direct comparisons (Figure 2). *mNG^SEC^3xFlag::AID::mpk-1* knock-in animals generated using the original SEC could not be maintained as homozygotes prior to SEC excision (Figure 2A), as expected based on essential functions for ERK/MPK-1. mNG protein in these knock-in animals localized uniformly throughout the cytoplasm and nucleus (Figure 2B). In contrast, *mNG::AID^NSEC^mpk-1* knock-in animals were viable as homozygotes (Figure 2A) and did not exhibit *mpk-1* loss-of-function phenotypes. In all cell types examined, the subcellular localization of mNG::AID^NSEC^MPK-1 protein (Figure 2B) resembled published knock-in strains (RASMUSSEN AND REINER 2021; GIBNEY *et al*. 2025) consistent with production of functional mNG::AID::MPK-1 protein prior to selectable marker excision in *mNG::AID^NSEC^mpk-1* animals. To verify that the NSEC design remained capable of self-excision, we heat shocked young L1 stage *mNG::AID^NSEC^mpk-1* worms and screened their progeny for individuals without the roller phenotype. We observed numerous F1 animals with wild-type movement and picked a single founder to establish an excised, homozygous *mNG::AID::mpk-1* line (Figure 2A). mNG::AID::MPK-1 protein localization was indistinguishable between the NSEC and excised lines (Figure 2B). The combination of biological functionality and correct subcellular localization prior to NSEC excision indicates that the NSEC does not interfere with the function of an essential N-terminally-tagged protein. To streamline use of this method by the *C. elegans* community, we developed a plasmid toolkit featuring NSEC backbones with five codon-optimized fluorescent proteins with and without an AID tag (Table 1).

**Table 1.** Plasmids for NSEC-based endogenous gene tagging

| Plasmid | Description | Purpose | Addgene ID |
| --- | --- | --- | --- |
| pTG618 | mTurquoise<sup>NSEC</sup>linker | endogenous gene tagging with mTurquoise2 | 237294 |
| pTG619 | mTurquoise::AID<sup>NSEC</sup>linker | endogenous gene tagging with mTurquoise2::AID | 237295 |
| pTG614 | GFP<sup>NSEC</sup>linker | endogenous gene tagging with GFP | 237290 |
| pTG615 | GFP::AID<sup>NSEC</sup>linker | endogenous gene tagging with GFP::AID | 237291 |
| pTG612 | mStayGold<sup>NSEC</sup>linker | endogenous gene tagging with mStayGold | 237387 |
| pTG613 | mStayGold::AID<sup>NSEC</sup>linker | endogenous gene tagging with mStayGold::AID | 237239 |
| pTG610 | mNeonGreen<sup>NSEC</sup>linker | endogenous gene tagging with mNeonGreen | 237385 |
| pTG611 | mNeonGreen::AID<sup>NSEC</sup>linker | endogenous gene tagging with mNeonGreen::AID | 237386 |
| pTG616 | mScarlet-I<sup>NSEC</sup>linker | endogenous gene tagging with mScarlet-I | 237292 |
| pTG617 | mScarlet-I::AID<sup>NSEC</sup>linker | endogenous gene tagging with mScarlet-I::AID | 237293 |

**Figure 2.**
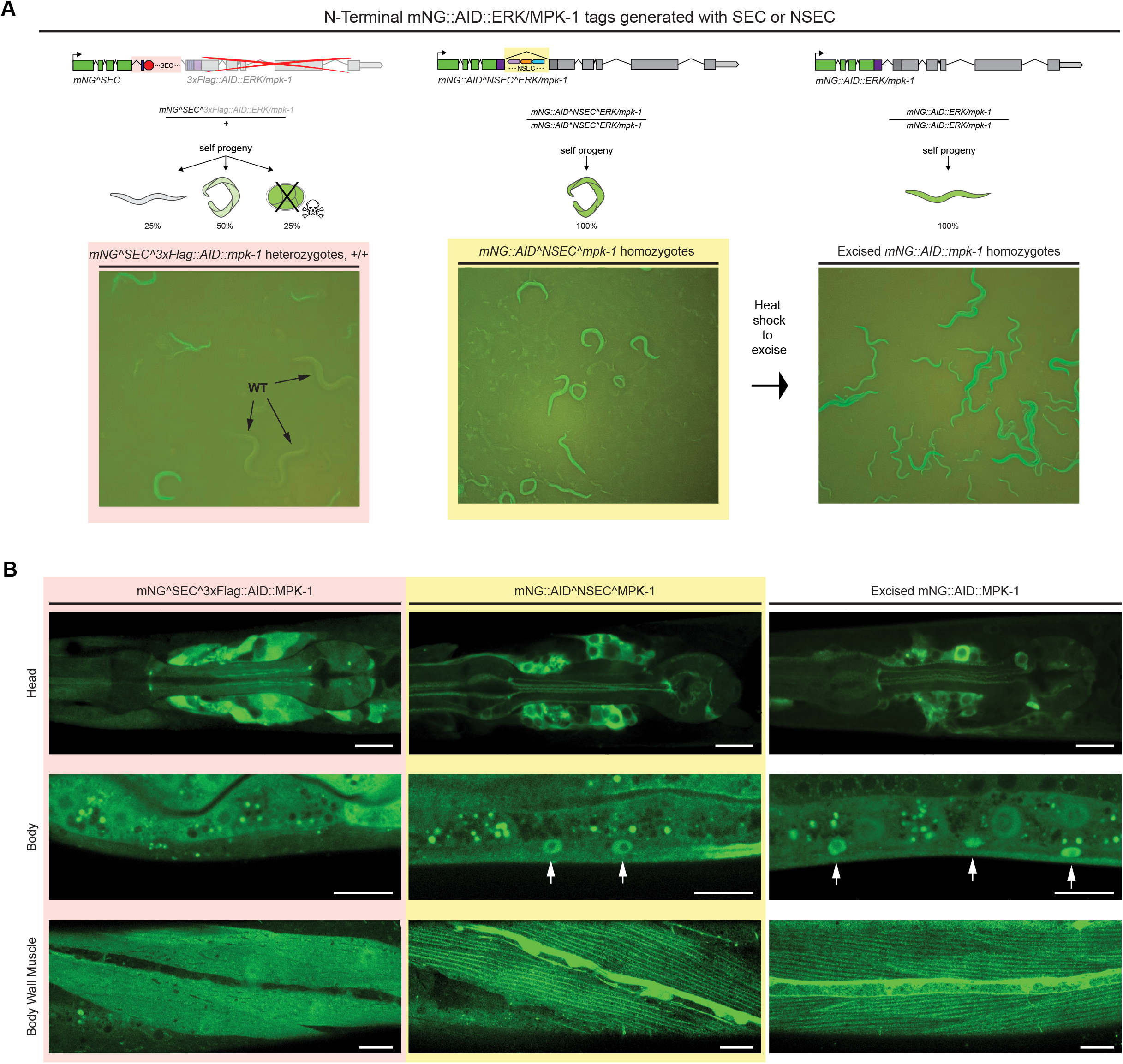
N-terminal NSEC knock-in does not interfere with endogenous ERK/MPK-1 function or localization.

## Conclusions

The NSEC provides a user-friendly approach to tag endogenous genes in any location without initially disrupting expression or function (Figure 3). Because N-terminal or internal tags are often required to preserve the functionality of tagged proteins, we expect the NSEC will streamline the process of generating endogenously tagged alleles for many genes. The ability to use a standard workflow for N-terminal, internal, and C-terminal tags without the need for genetic balancers simplifies the process of making knock-ins for essential genes and should facilitate endogenous tagging efforts in *C. elegans*. The ability to maintain homozygous NSEC lines for N-terminally tagged essential genes prior to selectable marker excision also eases strain maintenance and makes it possible to easily genotype subsequent crosses using the roller phenotype. Compared to methods that rely on screening for edited animals by PCR or fluorescence, the use of selectable markers makes it possible to recover knock-ins at sites with low insertion efficiency and/or for genes expressed at low levels with minimal hands-on effort. While we have only tested the NSEC for fluorescent protein-tagging, this method should allow for other types of genome edits and insertions by modifying the homologous repair templates using conveniently placed restriction sites (see Figure 1; Supplemental Information). For labs that are familiar with the existing SEC workflow (DICKINSON *et al*. 2015), using the NSEC should be seamless as the process remains identical except for the use of a different repair template plasmid. To facilitate adoption, we provided ten NSEC repair template backbones that meet a range of experimental needs. Beyond *C. elegans*, this strategy has potential applications in other model systems, providing a framework for scarless knock-ins that retain endogenous gene function during all steps of the process. By easing the burden of tagging essential genes, the NSEC will facilitate large-scale efforts to endogenously tag *C. elegans* genes and accelerate the types of experiments needed for a mechanistic understanding of fundamental biological processes *in vivo*.

**Figure 3.**
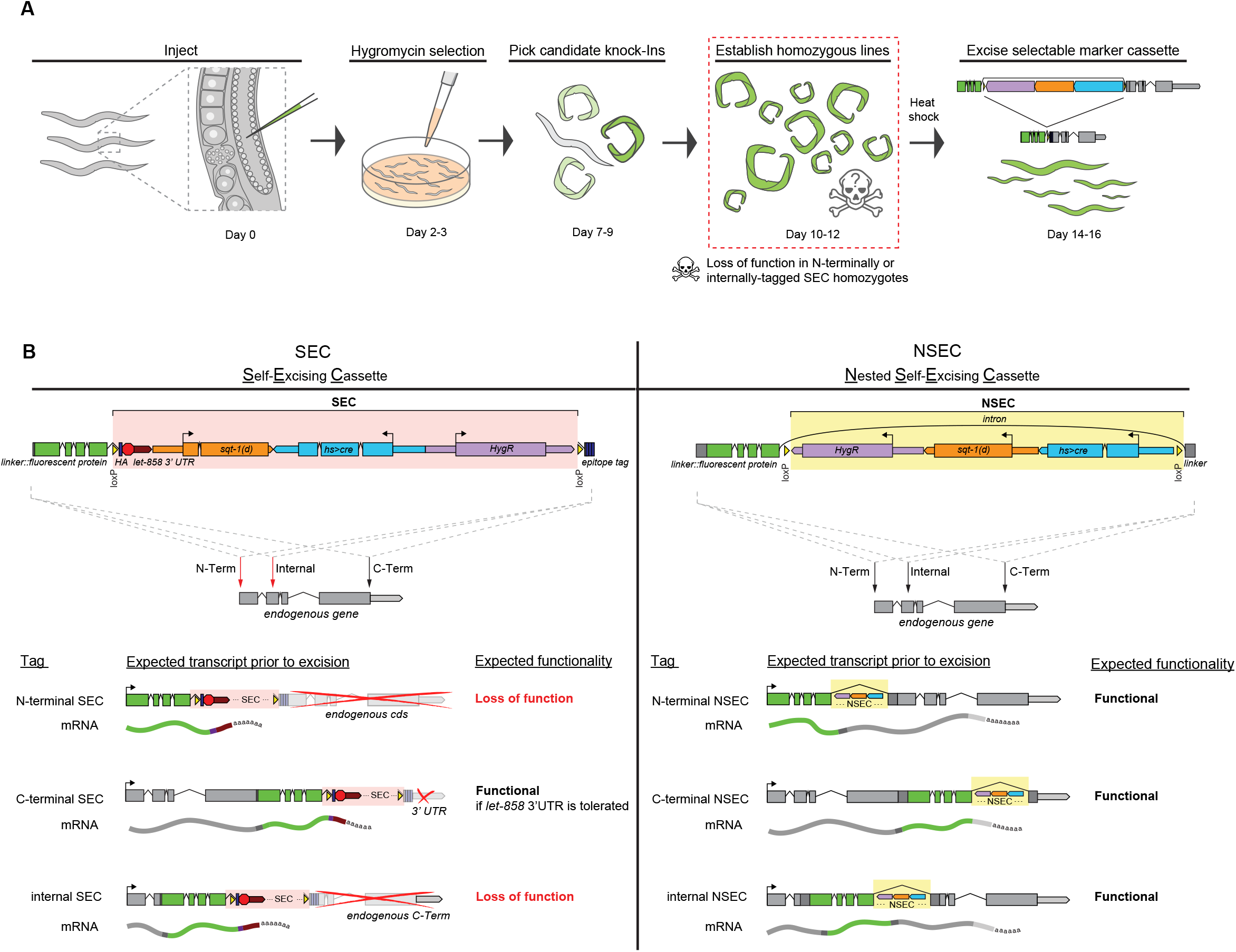
Workflow and expected outcomes for endogenous gene tagging with the SEC or NSEC strategies.

## Data and Materials Availability

NSEC plasmid backbones generated in this study are listed in Table 1 and have been deposited at Addgene. Additional materials are listed in Table S1 (Resources and Reagents). Annotated plasmid sequence files are provided in Supplemental File 1. Original image data are available upon request. For further inquiries or requests, please contact Ariel Pani at.

## Acknowledgements

This research was funded by National Institute of General Medical Sciences grant R35GM142880 (AMP) and Eunice Kennedy Schriver National Institute of Child Health and Human Development fellowship F31HD112152 (TVG). The wild-type N2 strain was provided by the CGC, which is funded by the NIH Office of Research Infrastructure Programs (P40 OD010440).

## Author Contributions

Conceptualization, TVG and AMP; methodology, TVG and AMP; investigation, TVG and AMP; data curation, TVG; writing – original draft, TVG; writing – review & editing AMP; visualization, TVG and AMP; project administration, AMP; funding acquisition, TVG and AMP.

## Materials and methods

### *C. elegans* maintenance

*C. elegans* knock-in strains were generated in a wild-type (N2) background. All *C. elegans* strains were maintained at room temperature (approximately 22°C) on Nematode Growth Media (NGM) plates seeded with *E. coli* OP50 as described in detail elsewhere (STIERNAGLE 2006).

### Plasmid design and molecular cloning

We identified potential intron splice acceptor sites using neural network-based prediction implemented through the NetGene2 - 2.42 server with *C. elegans* selected as the species (
https://services.healthtech.dtu.dk/services/NetGene2-2.42/; BRUNAK *et al*. 1991; HEBSGAARD *et al*. 1996). We then made synonymous substitutions to eliminate predicted splice acceptors within coding sequences without altering the encoded protein. For predicted splice acceptors in noncoding sequence, we either altered or added individual bases to eliminate the predicted splice acceptor function. Alterations made relative to the original SEC sequence are detailed in Supplemental Note 1. Including an intron in *cre* was essential to prevent bacterial *cre* expression and resulting Cre-lox recombination within the NSEC plasmids during cloning. The NSEC sequence was synthesized commercially (GENEWIZ, Azenta Life Sciences). To generate pTG610 and pTG611, we used Gibson assembly (New England Biolabs HiFi DNA Assembly) to add flanking intron, linker, and mNG or mNG::AID sequences. Because some fluorescent protein fusions require longer linker sequences than used in the original SEC to retain biological function (GIBNEY *et al*. 2025), we added new extended, flexible linker sequences. To generate additional repair templates with other fluorescent proteins, the mNG coding sequence in pTG610 or pTG611 was replaced with mTurquoise2, GFP, mStayGold, or mScarlet-I using Gibson assembly. Homologous repair template plasmids for mNG^SEC^3xFlag::AID::mpk-1 and mNG::AID^NSEC^mpk-1 were made from pUA77 (AGHAYEVA *et al*. 2021) or the precursor of pTG611, respectively. PCR fragments were amplified from existing plasmids or genomic DNA using Q5 Hi-Fidelity 2X Master Mix (New England Biolabs). Plasmids were transformed into DH5-alpha cells (New England Biolabs), and miniprepped with a PureLink HQ Mini Plasmid DNA Purification Kit (Invitrogen). Site-directed mutagenesis of pDD162 (DICKINSON *et al*. 2013) was used to generate the Cas9 + N-terminal *mpk-1* guide RNA plasmid pTG173. *mpk-1* was endogenously tagged immediately following the start codon of F43C1.2a using the guide RNA sequence 5’ TTCTTCTTGCAGATGGCCGA 3’, which is predicted to result in an N-terminal tag for MPK-1 isoform A and an internal tag for isoform B (STERNBERG *et al*. 2024). All plasmid sequences were confirmed with whole-plasmid nanopore sequencing (Plasmidsaurus).

### Strain construction

*C. elegans* knock-in strains using either SEC- or NSEC-based plasmid repair templates were generated by germline microinjection as described in detail elsewhere (DICKINSON *et al*. 2015; GIBNEY *et al*. 2023) (https://wormcas9hr.weebly.com/). Briefly, we injected a mix containing a repair template plasmid, Cas9 + guide RNA plasmid pTG173, and red fluorescent co-injection markers (DICKINSON *et al*. 2013) into the germlines of young adult hermaphrodites. Following microinjections, animals were moved to fresh, NGM plates seeded with *E. coli* OP50 (3-4 worms per plate). Plates were treated with hygromycin 2-3 days post-injection, prior to reproductive maturity of the F1 progeny. We screened for candidate knock-ins 7-9 days post-injection by looking for plates with numerous young, rolling animals that lacked the red fluorescent co-injection markers. Sixteen animals from each candidate plate were then picked onto individual plates without hygromycin. These plates were assessed three days later for homozygosity by screening for plates where 100% of progeny exhibited the roller phenotype. Because *mNG^SEC^3xFlag::AID::mpk-1* could not be maintained as a homozygote, we maintained this strain as heterozygotes by picking rolling animals with superficially normal development. The NSEC was excised from homozygous *mNG::AID^NSEC^mpk-1* (APL958) animals by heat shocking approximately 30 young L1 stage worms at 34°C for four hours. We then established the excised *mNG::AID::mpk-1* line (APL959) by picking progeny with wild-type movement four days after heat shock.

### Microscopy

Images in Figure 2A were taken using an Apple iPhone15 and Zeiss Axio Zoom V16 fluorescence microscope with a Plan NeoFluar Z 2.3x/0.57 objective. Images in Figure 2B were acquired using a Yokogawa CSU-X1 spinning disk confocal and Hamamatsu ORCA Fusion BT sCMOS camera mounted on a Nikon Ti2E inverted microscope stand. Spinning disk imaging was performed using an Apo TIRF 60X/1.49 NA oil immersion objective with 514 nm laser excitation, 445/514/594 dichroic mirror, and 545/40m emission filter. Larval worms were immobilized for live imaging in 0.3 mmol/L levamisole in M9 buffer and mounted on 3% (wt/vol) agarose pads. Strains were imaged within 30 minutes of mounting, and at least 50 worms were examined per strain. To compare protein localization between strains, animals were imaged at comparable developmental stages. Images were acquired using Nikon NIS Elements AR 5.42 software. Images were deconvolved using Nikon NIS Elements and adjusted for brightness and contrast using Fiji/ImageJ (SCHINDELIN *et al*. 2012). Figures were prepared using Adobe Illustrator 28.3.

## Figure and Table Legends

**Figure 1. Design of the Nested, Self-Excising selection Cassette (NSEC)**. (**A**) The NSEC consists of a hygromycin resistance gene, *sqt-1(d)* dominant phenotypic marker, and heat-shock inducible *cre* flanked by loxP sites and embedded within a synthetic intron. Cre-lox recombination can excise the NSEC, leaving behind a seamless genomic insertion. The mNG::AID^NSEC^linker plasmid is shown for illustrative purposes, but the design is similar for all versions described here. (**B**) Schematic of plasmid backbone pTG611 corresponding to the NSEC design in A. Locations of restriction sites for inserting homology arms and making other repair template modifications are highlighted. See Supplemental Information for plasmid sequences and annotations.

**Figure 2. N-terminal NSEC knock-in does not interfere with endogenous ERK/MPK-1 function or localization**. (**A**) Architecture and functionality of N-terminal SEC and NSEC knock-ins tagging the essential gene *mpk-*1. An *mNG^SEC^3xFlag::AID::mpk-1* knock-in disrupts function of *mpk-1* due to the design of the original SEC, which includes a strong transcriptional terminator at the 5’ end of the SEC. *mNG^SEC^3xFlag::AID::mpk-1* animals could only be maintained as heterozygotes, which segregated wild-type animals, fertile heterozygous knock-in animals, and sterile homozygous knock-in animals with loss of MPK-1 function (maternally rescued homozygotes are viable but sterile). In contrast, *mNG::AID^NSEC^mpk-1* knock-in animals were viable as homozygotes and did not exhibit loss-of-function phenotypes. The NSEC can be self-excised by heat shock to remove the selectable marker cassette and restore a wild-type movement phenotype. (**B**) Expression and subcellular localization of SEC- and NSEC-tagged endogenous MPK-1. *mNG^SEC^3xFlag::AID::mpk-1* knock-in animals expressed only mNG, which localized uniformly throughout both the cytosol and nuclei of cells that express *mpk-1*. In contrast, subcellular localization of mNG::AID^NSEC^MPK-1 was indistinguishable from excised mNG::AID::MPK-1. The tagged protein in both NSEC and excised strains is excluded from the nuclei of multiple cells in the head but enriched in nuclei (arrows) in other cell types including vulval precursors and muscles where ERK signaling is active. mNG::AID^NSEC^MPK-1 and excised mNG::AID::MPK-1 protein also exhibited a striated localization pattern in body wall muscles. Scale bars = 10µm.

**Figure 3. Workflow and expected outcomes for endogenous gene tagging with the SEC or NSEC strategies**. (**A**) The SEC and NSEC strategies use identical workflows to generate endogenous knock-ins with the exception that the SEC can interfere with endogenous gene function prior to excision. (**B**) Schematic diagrams of SEC and NSEC architectures and expected outcomes for endogenous knock-ins in different locations within a gene. The SEC includes a *let-858* 3’UTR with a strong transcriptional terminator that is appended to the fluorescent protein prior to excision. N-terminal and internal tags using the SEC are expected to result in loss of endogenous gene function prior to SEC excision. The NSEC is embedded entirely within a synthetic intron and includes no splice acceptors, stop codons, or transcriptional terminators in the same orientation as the gene of interest. NSEC knock-ins in any location within a gene are expected to retain endogenous gene expression and function prior to NSEC excision.

**Table 1. Plasmids for NSEC-based endogenous gene tagging**. Names, descriptions, and Addgene accession numbers for NSEC plasmids for gene tagging with codon-optimized mTurquoise2, GFP, mStayGold, mNeonGreen, or mScarlet-I.

## Supplemental Information

**Supplemental Note 1**. (**A**) Modifications in the NSEC coding and regulatory sequences relative to the SEC. (**B**) Annotated sequence for pTG611 (mNG::AID^NSEC^linker).

**Table S1. Resources and Reagents**.

**Supplemental File 1. NSEC plasmid sequence files**. Single .zip file containing sequences for NSEC backbone plasmids and mNG::AID^NSEC^mpk-1 homologous repair template plasmid. Figshare link: https://doi.org/10.6084/m9.figshare.28893263.v1

## Supplemental Information including

Supplemental Note 1 – Annotated NSEC sequence features

Table S1 – Resources and Reagents

Supplemental File 1 – Plasmid sequences (.zip)

## Supplemental Note 1

### (A) Nucleotide substitutions in the NSEC coding and regulatory sequences relative to the SEC

Sequences are shown in the orientation present in the NSEC. Nucleotides that were edited to eliminate sense strand splice acceptors or restriction sites are shown in bold.

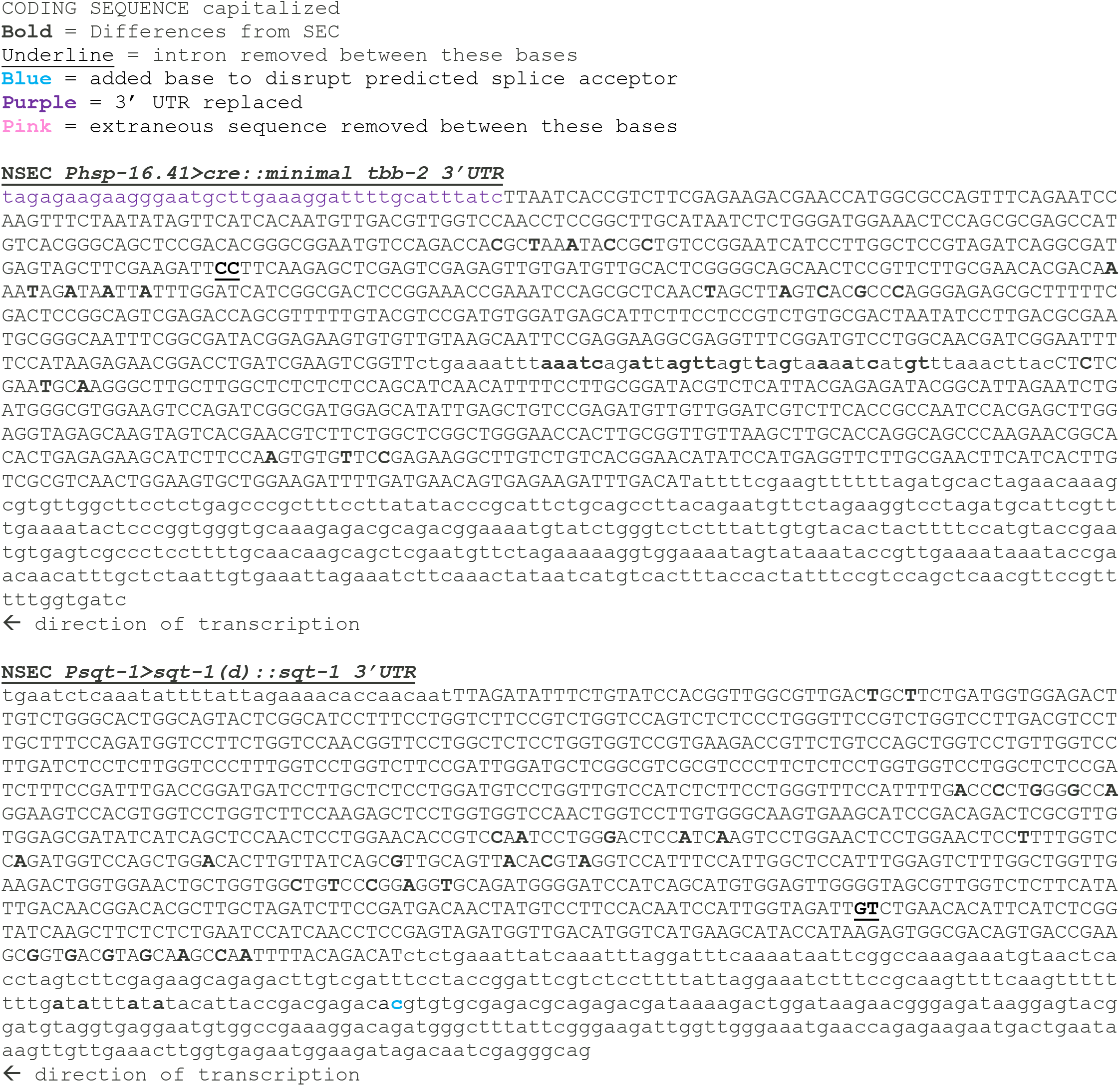
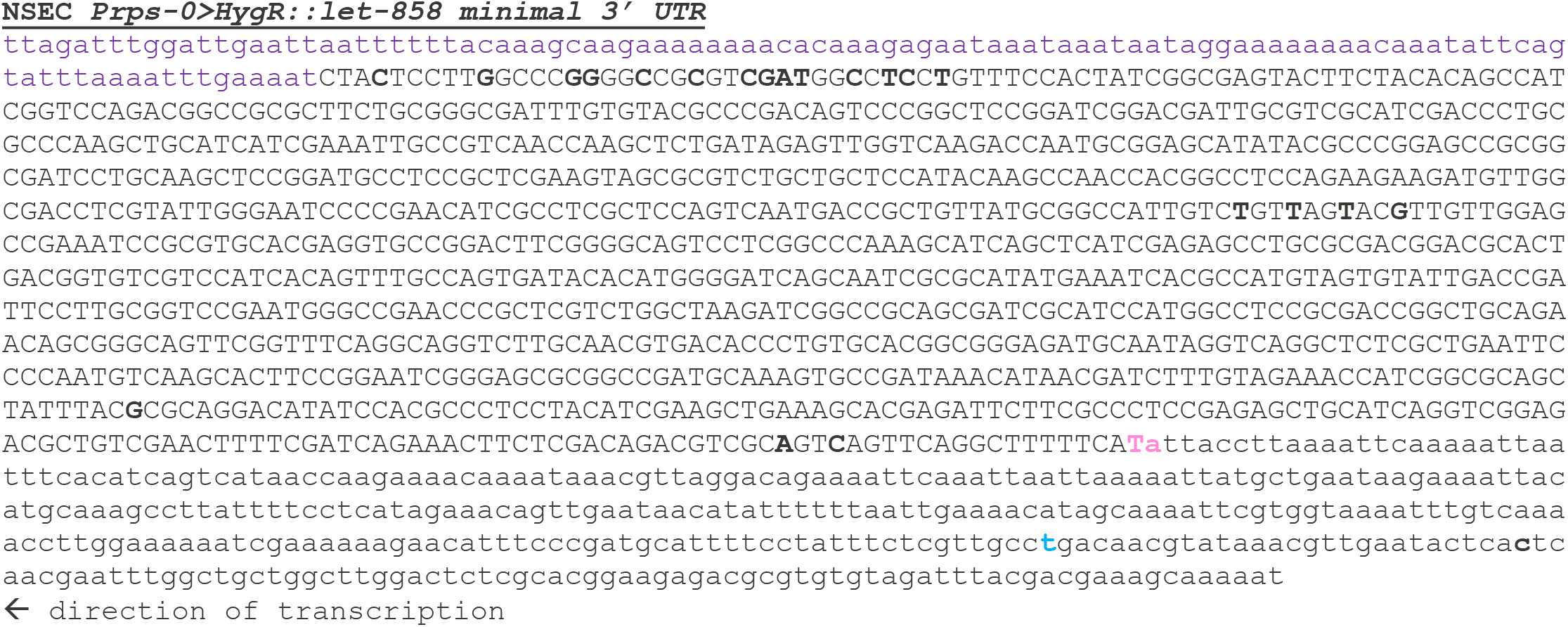

### (B) Annotated sequence for pTG611 (mNG::AID^NSEC^linker)

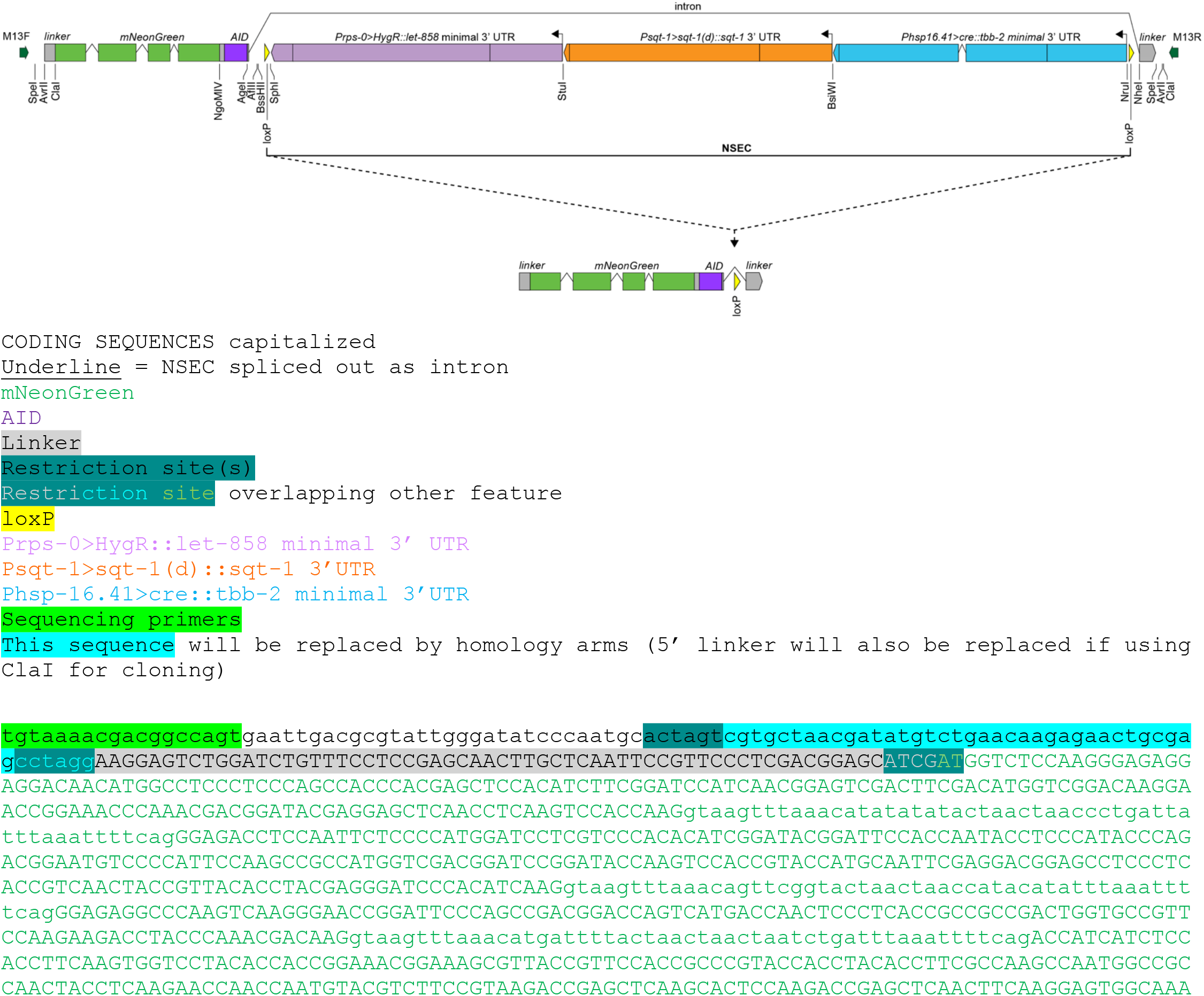
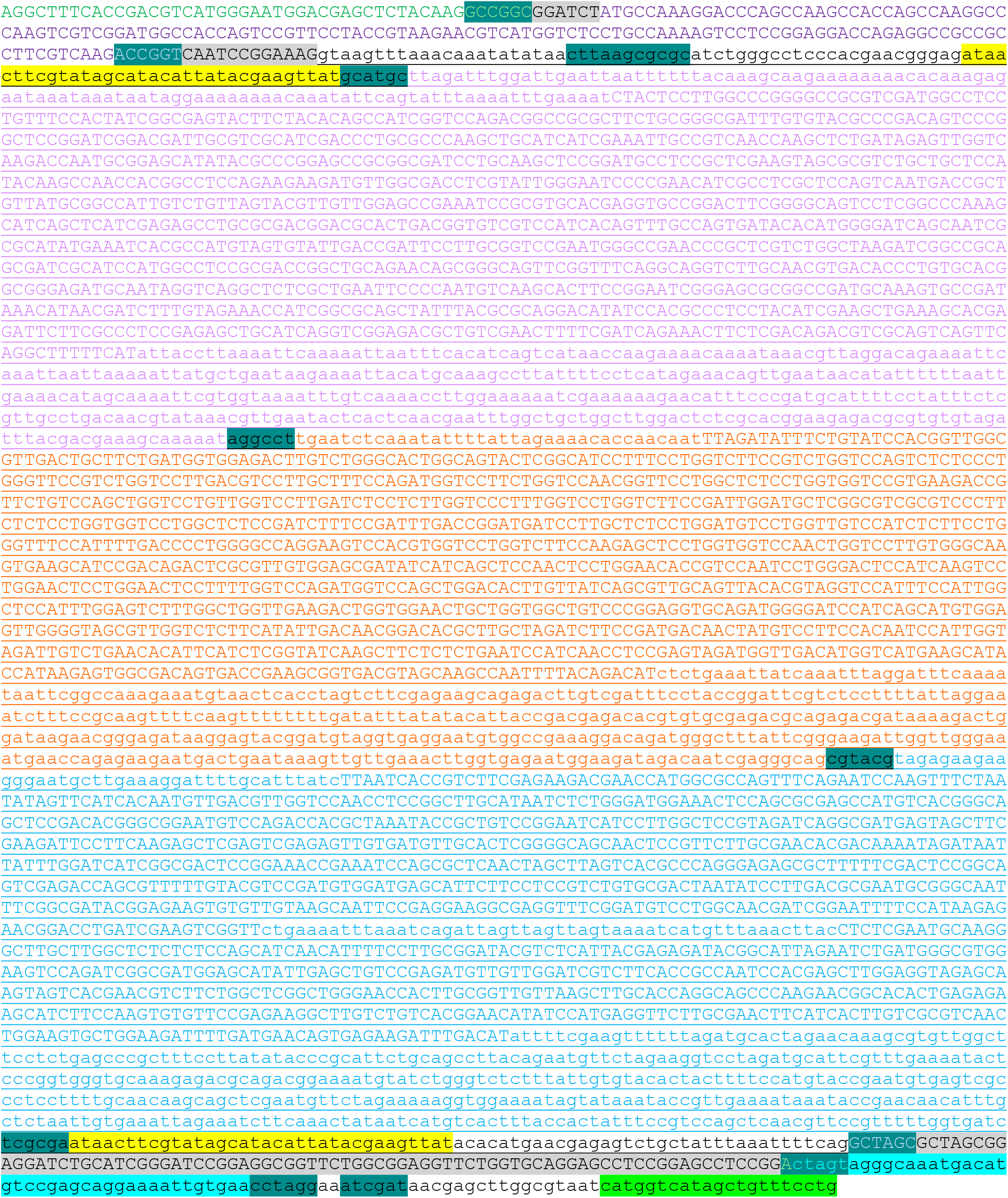

**Table S1.** Resources and Reagents

| Resource or Reagent | Source | Identifier |
| --- | --- | --- |
| ***C. elegans* strains** |  |  |
| wild-type | CGC | N2 |
| mpk-1(ljf105[mNG<sup>^</sup>SEC<sup>^</sup>3xFlag::AID::mpk-1]) III* | This paper | APL934 |
| mpk-1(ljf106[mNG::AID<sup>^</sup>NSEC<sup>^</sup>mpk-1]) III | This paper | APL958 |
| mpk-1(ljf107[mNG::AID::mpk-1]) III | This paper | APL959 |
| ***Plasmids*** |  |  |
| Peft-3>Cas9 + PU6>ttTi5605 sgRNA | Dickinson *et al.*, 2013 | pDD122; Addgene 47550 |
| Peft-3>Cas9 + PU6>mpk-1 N-terminus sgRNA | Gibney *et al.*, 2025 | pTG173 |
| mNG<sup>^</sup>SEC<sup>^</sup>3xFlag::AID::mpk-1 | Gibney *et al.*, 2025 | pTG174 |
| mNG::AID<sup>^</sup>NSEC<sup>^</sup>mpk-1 | This paper | pTG607 |
| mNeonGreen<sup>^</sup>NSEC<sup>^</sup>linker | This paper | pTG610; Addgene 237385 |
| mNeonGreen::AID<sup>^</sup>NSEC<sup>^</sup>linker | This paper | pTG611; Addgene 237386 |
| mStayGold<sup>^</sup>NSEC<sup>^</sup>linker | This paper | pTG612; Addgene 237387 |
| mStayGold::AID<sup>^</sup>NSEC<sup>^</sup>linker | This paper | pTG613; Addgene 237239 |
| GFP<sup>^</sup>NSEC<sup>^</sup>linker | This paper | pTG614; Addgene 237290 |
| GFP::AID<sup>^</sup>NSEC<sup>^</sup>linker | This paper | pTG615; Addgene 237291 |
| mScarlet-I<sup>^</sup>NSEC<sup>^</sup>linker | This paper | pTG616; Addgene 237292 |
| mScarlet-I::AID<sup>^</sup>NSEC<sup>^</sup>linker | This paper | pTG617; Addgene 237293 |
| mTurquoise<sup>^</sup>NSEC<sup>^</sup>linker | This paper | pTG618; Addgene 237294 |
| mTurquoise::AID<sup>^</sup>NSEC<sup>^</sup>linker | This paper | pTG619; Addgene 237295 |
| ccdB mNG<sup>^</sup>SEC<sup>^</sup>3xFlag::AID ccdB | Aghayeva *et al.*, 2021 | pUA77 |
| ***Key reagents*** |  |  |
| Hygromycin B (50 mg/mL) (Gibco) | Fisher Scientific | 10-687-010 |
| PureLink HQ Mini Plasmid DNA Purification Kit | Invitrogen | K210001 |
| Q5 High-Fidelity 2X Master Mix | New England Biolabs | M0492L |
| Q5 Site-Directed Mutagenesis Kit | New England Biolabs | E0554S |
| NEBuilder HiFi DNA Assembly Cloning Kit | New England Biolabs | E5520S |
| NEB 5-alpha Competent *E. coli* (High Efficiency) | New England Biolabs | C2987H |
| ***Software and algorithms*** |  |  |
| Fiji/ImageJ2 (Version: 2.14.0/1.54f) | Schindelin *et al.*, 2012 |  |
| Adobe Illustrator CC (Version 28.3) | Adobe Systems |  |
| Nikon NIS-Elements (Versions AR 5.42.06 and AR 5.42.04) | Nikon Instruments Inc. |  |
| NetGene 2 - 2.42 (DTU Health Technology) | Brunak *et al.*, 1991; Hebsgaard *et al.*, 1996 |  |

\* APL934 can only be maintained as a heterozygote. Maternally rescued homozygotes are sterile.

**Supplemental File 1. NSEC plasmid sequence files**. Single .zip file containing sequences for NSEC backbone plasmids and mNG::AID^NSEC^mpk-1 homologous repair template plasmid. Figshare: https://doi.org/10.6084/m9.figshare.28893263.v1

